# Multispecies interactions and the community context of the evolution of virulence

**DOI:** 10.1101/2024.03.29.587402

**Authors:** Claire Evensen, Andy White, Mike Boots

## Abstract

In nature, host-parasite/pathogen relationships are embedded in a network of ecological interactions that have the potential to shape the evolutionary trajectories of shared pathogens. Understanding this community context of infectious disease evolution is important for wildlife, agricultural, and human systems alike – illustrated, for example, by the increasing risk of zoonotic disease emergence. We introduce an eco-evolutionary model that examines ecological feedbacks across a range of host-host interactions. Specifically, we analyze a model of the evolution of virulence of a pathogen infecting hosts who themselves exhibit competitive, mutualistic, or exploitative relationships. We find that pathogen specialism is necessary for inter-host interactions to impact parasite evolution. An important general result is that increasing competition between hosts leads to higher shared pathogen virulence, while increasing mutualism leads to lower virulence. Across a range of scenarios, the nature of pathogen specialization is critical to the outcome – for instance, if hosts only differ in initial susceptibility to infection, there is no impact of host-host interactions on virulence evolution. In contrast, specialization in terms of onward transmission, host tolerance, or intra-host pathogen growth rate critically impact the evolution of virulence. For example, stronger specialism in transmission selects for lower virulence, while stronger specialism in tolerance and growth rate selects for higher virulence. Our work provides testable hypotheses for multi-host disease systems, predicts how changing interaction networks may impact the evolution of virulence, and broadly demonstrates the importance of looking beyond pairwise relationships to understand evolution in realistic natural contexts.

## Introduction

Parasitic lifestyles are ubiquitous and the wider importance of infectious disease to human health, agriculture, and natural systems is clear.^1–4^ As a consequence there is a large body of theoretical work focused on understanding the evolution of pathogen virulence.^5–8^ Consistent with their pervasiveness, host-parasite interactions in nature have not evolved in isolation, but rather in complex webs of species interactions. Multi-host parasites, or those that can infect and transmit onward from more than one host, are the norm, not the exception^9^ – they dominate infections in wildlife communities^10, 11^ and of course many human pathogens are zoonotic.^12, 13^ However, the evolutionary consequences of these ubiquitous multi-host interactions on pathogen evolution have only rarely been considered,^14^ and there is a lack of theory focused on how host interactions will interact with epidemiological processes to affect virulence evolution.^5, 15, 16^.

It has been shown that multi-host parasite fitness typically varies across hosts^14^ with theory proposing that parasites should evolve optimal virulence on their highest quality hosts.^17^ The most obvious (though often difficult to determine) metrics for host quality are tied to the mechanisms of infection and transmission. If a pathogen performs well on one host and poorly on another, the difference could be attributed to a greater susceptibility of one host to initial infection, a higher intra-host pathogen growth rate, a greater efficiency of production of pathogen propagules for onward transmission, a lower death rate due to infection (i.e. tolerance), or a mixture of these mechanisms.^18–21^ The precise reasons for fitness variation and subsequent evolutionary consequences are not so easily determined, however, as host quality is a product of not only underlying biology, but also host population interactions.^22^ For example, a host that is more susceptible to a parasite may have a reduced population size and this will have impacts both directly on parasite numbers and the population size of other hosts through interspecific interactions. We therefore need to consider the tangled web of interactions in which a parasite’s hosts are embedded since host population dynamics are readily impacted by non-parasitic ecological interactions, as has been demonstrated in micro- and macroorganism systems.^23–26^ It is clear that such complex eco-evolutionary feedbacks may have important implications to the evolution of virulence,^27^ but we have little understanding of how interactions between different hosts in multi-host infectious disease systems impact parasite evolution.

To date, there is a large body of theoretical and empirical work on the disease ecology and evolution in pairwise one host-one parasite relationships.^5, 28, 29^ Multi-host systems have been explored in the context of population dynamics and epidemiology, where the consequences of varying inter-vs intra-host disease transmission and the trade-offs between virulence in different hosts have been considered.^17, 30, 31^ Interspecies transmission and intrinsic host mortality have been identified as key determinants of multi-host parasite evolution,^14, 17, 32^ but other forms of interspecies interactions have been ignored. Given the importance of disease spillover in multi-host, multi-parasite systems,^33–35^ the question of which host communities may favor the evolution of high parasite virulence is critical. Recent work has begun to explore parasite evolution in more complex ecological contexts, such as hyperparasitism^36, 37^ and multi-parasite assemblages,^38, 39^ and has called for further exploration of the consequences of inter-host interactions on shared parasites,^40^ but our theoretical understanding of the evolution of parasites in a realistic community context remains limited.

Here, our goal is to develop a theoretical foundation for predicting evolutionary trajectories of multi-host parasites, over a range of different interspecific host interactions. We develop a two-host, one-parasite model to investigate our central question of how parasite virulence evolves under host-host interactions ranging from antagonism and mutualism to exploitation, when parasites vary in their degree of host specialism. To paint a more complete picture of how these evolutionary trajectories are influenced by the degree to which a parasite specializes on one of several hosts, we explore the interaction of host-host relationships with distinct mechanisms that give rise to variation in parasite performance across their hosts. We show that the inter-host interaction is critical to the evolution of specialist parasites.

## Methods

### Two host, one parasite model framework

We modify a standard SIS model for one host and one parasite, extending to two hosts and one shared parasite. Hosts are capable of both intra-host and inter-host disease transmission. The dynamics of susceptible, *S*_*i*_, and infected, *I*_*i*_, hosts of type *i* = 1 and 2 are described by the following set of equations:

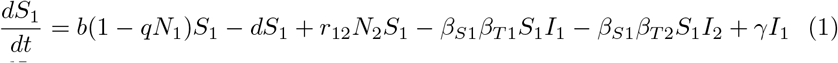

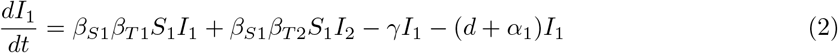

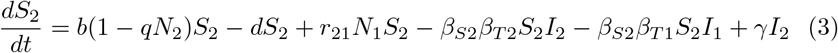

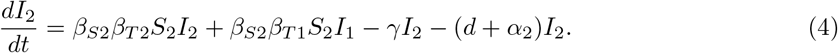

The parameters in the model are defined in Table 1 and a schematic of the model system is shown in Figure 1. To focus on the impact of host-host interactions, we assume that the following parameters are the same in both hosts: birth (*b*_1_ = *b*_2_ = *b*), death (*d*_1_ = *d*_2_ = *d*), crowding (*q*_1_ = *q*_2_ = *q*) and recovery (*γ*_1_ = *γ*_2_ = *γ*). The terms *α*_*i*_ represent the disease-induced mortality rate, or virulence, for host type *i*. We draw attention to the various *β* terms utilized. In order to explore different mechanisms by which parasites could vary in performance on each host (explained further in Results below), we expand the ‘typical’ *β* found in SI models to the product *β* = *β*_*Si*_*β*_*Tj*_, where *β*_*Si*_ is controlled by the host and represents a scaling factor that determines the susceptibility of host *i. β*_*Tj*_ is the transmission component controlled by the parasite. We note that work on multihost parasites often includes a *β*_*ij*_ term that reflects differences between inter- and intra-host contact rates. In our work we set equal inter- and intra-host contact rates, as we sought to isolate the impacts of host-host interactions. Such a term thus becomes another scaling factor that is incorporated (identically) into the *β*_*Si*_ term, and for clarity we do not explicitly include it in the model. Previous work often assumes a trade-off between virulence and transmission, in that *β* = *f* (*α*). In our breakdown of *β* into two parts, only *β*_*Tj*_ maintains a functional relationship with another parasite-related parameter. Furthermore, we decouple the traditional fixed relationship between *α* and *β* and explicitly incorporate *ϵ*, the parasite intra-host growth rate, setting the positive trade-off relationships between parasite growth rate, transmission, and host mortality accordingly:^28, 41^ *α*_*i*_ = *f* (*ϵ*_*i*_) and 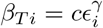, where *c* is a constant and *γ* < 1. For the former, in many cases this is a direct relationship of *α*_*i*_ = *ϵ*_*i*_ (thus preserving the same concave *β* = *f* (*α*) trade-off shape in previous work^42^), but see Figure 2 for the cases when differential parasite performance between the two hosts requires us to deviate from this precedent.

**Table 1:** Descriptions of model parameters.

| Symbol | Meaning |
| --- | --- |
| $N_i$ | total hosts of type $i$ |
| $S_i$ | susceptible hosts of type $i$ |
| $I_i$ | infected hosts of type $i$ |
| $b_i$ | natural birth rate of host type $i$ |
| $d_i$ | natural death rate of host type $i$ |
| $q_i$ | susceptibility to intraspecific crowding |
| $\epsilon_i$ | parasite growth rate in host type $i$ |
| $\alpha_i$ | parasite virulence (increased host mortality) |
| $\beta_{Si}$ | susceptibility of host type $i$ |
| $\beta_{Ti}$ | onward transmission from host type $i$ |
| $\gamma_i$ | recovery rate of host type $i$ |
| $r_{ij}$ | impact of host $j$ on host $i$ (interaction) |

**Figure 1:**
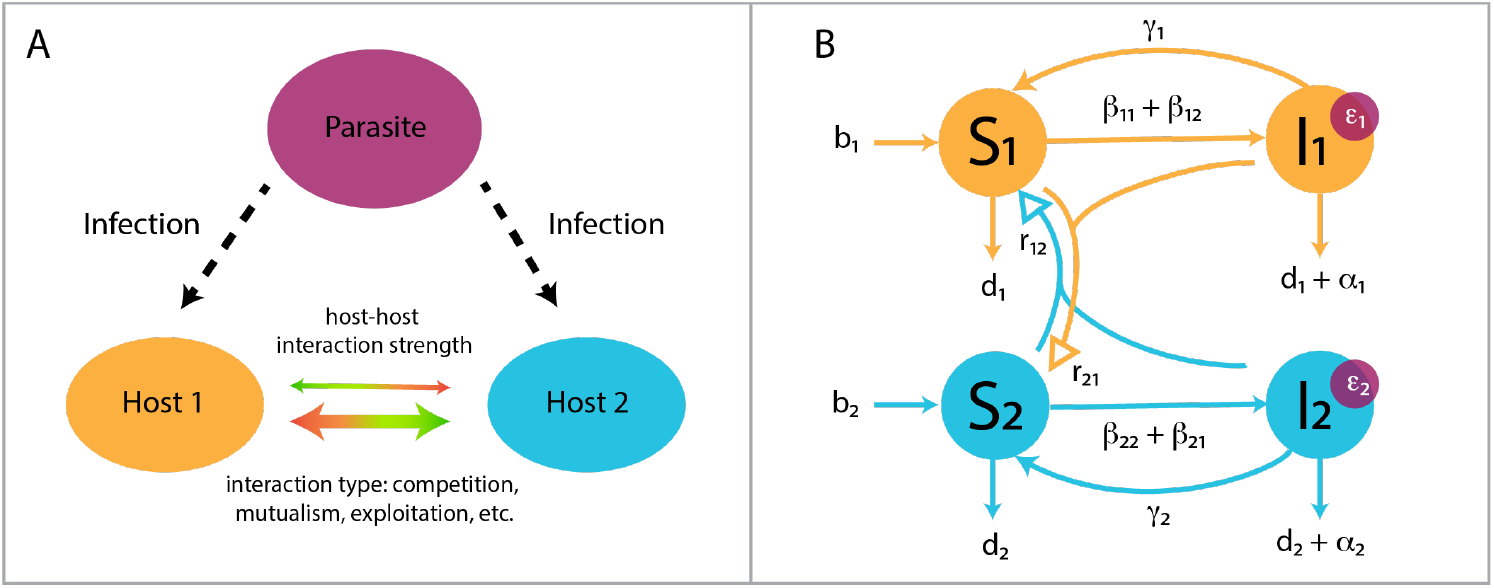
A schematic diagram of the model of the two-host, one parasite system represented in equations (1-4). Panel A highlights the key interactions we model: the simultaneous inter- and intra-host type infection dynamics of a shared parasite, and the ecological feedback generated by interhost interactions of varying strengths and types. Panel B shows the formal flowchart diagram for the model. Arrows with solid arrowheads indicate movement of individual hosts between susceptible (S) and infected (I) compartments through the processes of birth, death, infection, and recovery. Arrows with empty arrowheads do not indicate movement between host classes, but rather the impact (*r*_*ij*_) of host type *j* on the susceptible class of host type *i. β*_*ij*_ terms represent the composite transmission coefficient *β*_*Si*_*β*_*Tj*_; *b*_*i*_ and *d*_*i*_ are the natural birth and death rates of host type *i*, respectively; *α*_*i*_ and *γ*_*i*_ are the disease-induced death rate and recovery rate in host type *i*, respectively; *ϵ*_*i*_ is the parasite growth rate in host type *i*.

**Figure 2:**
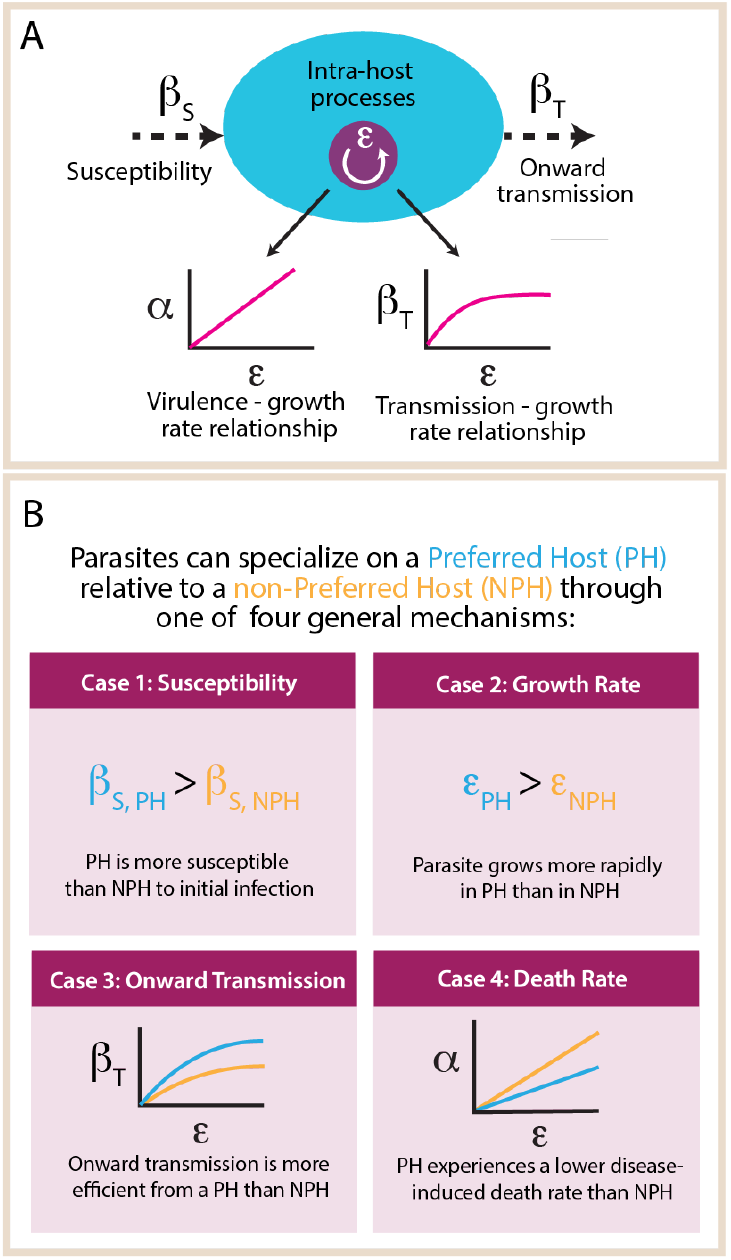
Mathematical representations of various mechanisms of parasite specialism. Panel A illustrates how we can break down the biology encompassed by a standard *β* transmission term into separate processes: a contact rate between hosts (not shown), susceptibility to an initial infection after contact (*β*_*S*_), intrahost parasite growth and replication (*ϵ*), and finally onward transmission of parasite propagules (*β*_*T*_). Panel B illustrates how we can use this detail to mathematically represent four different ways in which a parasite may specialize on one of its two hosts. The curves depicted in Cases 3 and 4 are predicated on standard assumptions that more aggressive parasite growth translates to greater harm done to the host (higher virulence *α*), and that the relationship between growth rate/virulence and onward transmission is saturating.

### Analytical and numerical techniques

We use the techniques of adaptive dynamics^43–46^ to determine the evolutionary stable level of virulence. These techniques assume that rare individuals arise from a homogeneous resident population via small mutations away from that resident strategy. The success of the rare mutant depends on its invasion fitness in the resident population – i.e. a positive initial growth rate for the mutant type in the resident population means the mutant strategy could, over time, coexist with or displace the resident strategy. As a consequence of our choice of trade-off, the population will evolve, through a process of sequential mutation, along its local fitness gradient until it reaches an evolutionarily stable strategy.

To represent the dynamics of hosts infected with the mutant parasites strain *I*_*iM*_ we add two equations to the system of Equations 1-4 as follows:

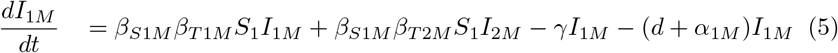

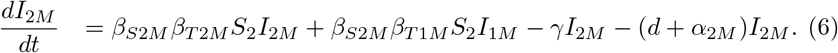

Therefore hosts can be infected with the resident or mutant parasite strain (though not both simultaneously). The full resident and mutant equations are shown in the supplementary information (Section S1).

To determine when invasion will occur, we analyze the eigenvalues of the Jacobian of the resident and mutant systems. In the absence of the mutant parasite we assume the resident system is at a stable equilibrium, (*Ŝ*_1_, *Î*_1_, *Ŝ*_2_, *Î*_2_). To examine whether the mutant can invade we determine the largest eigenvalue of the Jacobian at (*Ŝ*_1_, *Î*_1_, *Ŝ*_2_, *Î*_2_, 0, 0). For mutant invasion to occur, the resident-only equilibrium must be unstable, so we seek conditions where the system has a positive, real eigenvalue. It can be shown (see Section S1) that the mutant parasite strain can invade if the following condition holds:

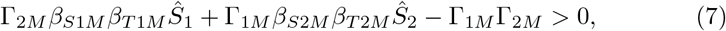

where Γ_1*M*_ = (*d* + *γ* + *α*_1*M*_) and Γ_2*M*_ = (*d* + *γ* + *α*_2*M*_). We use this fitness expression, with numerically determined values for the resident host populations (*Ŝ*_1_, *Ŝ*_2_) to construct pairwise invasion plots (PIPs)^43^(such as those in Figure 3) that allow us to determine the evolutionarily stable (ES) parasite virulence in a range of host-host interaction contexts. Each point in Figures 4-6 represents the ES virulence (determined from a PIP) for a particular parasite strategy and host-host interaction.

**Figure 3:**
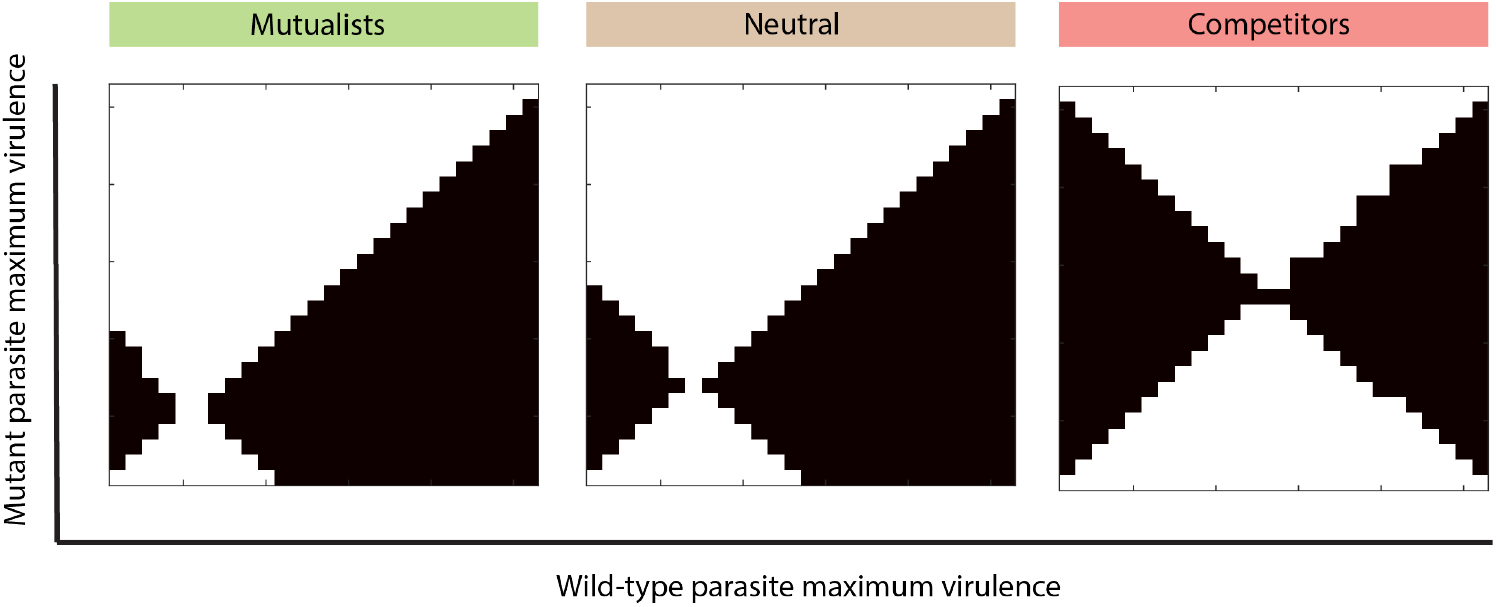
Pairwise invasibility plots that show the evolutionarily stable strategy for parasite virulence (ES virulence) on the backdrop of three different relationships between its two hosts: mutualists, neutral (no interaction beyond parasite transmission), and competitors. The points are continuously stable. Black regions show a pair of resident and mutant parasite strategies in which the mutant will be able to invade the wild-type resident population, and white regions show a pair in which the resident parasite will be able to outcompete the mutant. These analyses were performed for the case in which the parasite exhibits a higher growth rate in its preferred host than its non-preferred host (Case 2 specialism). The maximum virulence is thus the virulence realized on the preferred host, due to the relationship between growth rate and virulence described in the Methods section in the main text. Parameter values for all simulations can be found in Section S3.

**Figure 4:**
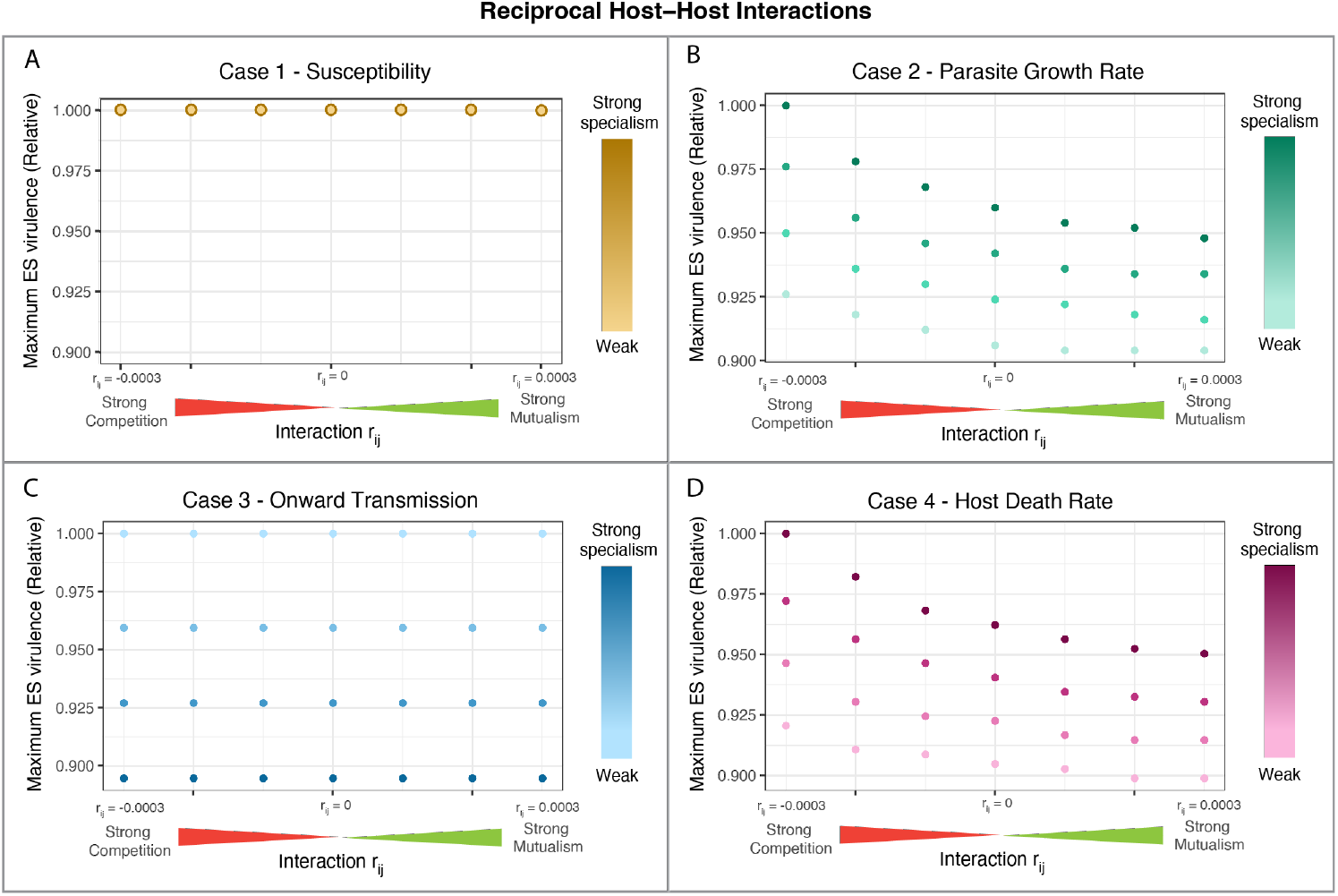
ES virulence for parasites of two reciprocally interacting hosts. In all the interactions tested here the two host interaction parameters were equal: *r*_*ij*_ = *r*_*ji*_. Reciprocal host interactions ranged from strong competition (*r*_*ij*_ = *r*_*ji*_ = −0.0003) to strong mutualism (*r*_*ij*_ = *r*_*ji*_ = 0.0003). We tested the range of host interactions on the backdrop of four different mechanisms of parasite specialism (Cases 1-4, panels A-D, respectively). In addition, four degrees of parasite specialism (ranging from strong to weak) were tested for each mechanism – strong specialism meant there was a large difference in parasite performance on each host type. The maximum ES virulence refers to the greater realized virulence, when realized virulence differs between host types (as occurs for Case 2 and 4 specialism). Because we do not compare magnitudes of ES virulence between cases, only within cases, we plot relative ES virulence (scaled from the highest value for a given specialism case). For the Case 1 plot, the degree of parasite specialism did not impact the ES virulence thus each point actually reflects four that are superimposed. For the exact parameter values for the degree of specialism, see Section S3.

## Results

We began by testing the simplest system: two hosts with identical life history parameters (birth, natural death, crowding) and identical disease burdens (specifically, identical values of *ϵ*, 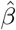, *α*), but with varying ecological relationships to one another. We find that shared parasite systems in which the parasite is a pure generalist (defined here as a parasite that has identical infection and disease characteristics in both hosts) do not yield different ES virulence values, even when the sign and magnitude of host-host interactions are varied. We therefore have a very general result that the nature of the interactions between hosts in multi-host parasite system has no impact on the evolution of a pure generalist parasite. We show an analytical proof of this result in the supplementary information (see Section S2).

We now include host specialism in the parasite, which reflects the observations in natural systems of multi-host parasites unevenly impacting their hosts – squirrelpox virus (SQPV) in grey and red squirrels,^47^ canine distemper virus (CDV) in a range of carnivores,^48^ and parasitic nematodes in *Drosophila*^21^ are several examples. Our definition of host types here can also allow for subdivision at a smaller level than species – even different host genotypes (with corresponding differences in parasite resistance or infection-induced mortality) could be considered distinct types.^49^ For the remainder of the paper, we will use specialism to refer to differential performance on hosts (not whether a parasite can infect a host or not).

In Figure 2, we outline four different potential mechanisms of parasite specialism and how they are modeled in our system. We distinguish between a preferred host (PH) and non-preferred host (NPH) based on which host the parasite would be described as performing “better”. Importantly, we note that this does not denote a deliberate host choice on the part of the parasite.

In Case 1, the parasite’s PH is more susceptible to infection than its NPH. In Case 2, the parasite’s PH sustains a more rapid parasite growth rate (and thus a higher virulence and rate of onward transmission, due to the dependence of the latter two on the former). In Case 3, for a given intra-host growth rate, the parasite transmits onward more effectively from its PH. In Case 4, for a given intra-host growth rate, the parasite causes less damage to its PH.

For each of the four specialism cases, we determined ES virulence for a range of host-host relationships, which we break down into two categories in the following sections: reciprocal and non-reciprocal interactions. Reciprocal interactions are defined as those with matching signs, therefore falling along a sliding scale of mutualism +/+ to competition −/−. Non-reciprocal interactions are defined as those with opposite signs, in that there is an antagonist (exploiter) that benefits and a subject of antagonism (exploitee) that suffers in the presence of the other host.

### Reciprocal interactions

We began by analyzing reciprocal host-host interactions of equal strength. When host-host interactions shifted ES virulence, they consistently followed a pattern of stronger mutualism leading to lower parasite virulence, and stronger competition leading to higher parasite virulence. Importantly, the four mechanisms of specialism did not lead to identical shifts in ES virulence. The cases in which the parasite reduced excess death in the PH and in which it grew more rapidly in the PH both led to significant shifts in virulence as interactions changed from competitive to mutualistic scenarios (Figure 4B, D), with virulence evolving to the highest levels under strongly competitive interactions and decreasing as the interaction progressed towards mutualistic interactions.

The case in which susceptibility was the only distinguishing factor between hosts did not show any shifts in ES virulence as host interactions changed, nor did the case in which onward transmission was more efficient in the PH (Figure 4A, C). The mathematical underpinnings of these two instances of independence between host-host relationship and pathogen virulence did differ. When the pathogen’s PH was more susceptible to infection (Case 1), the equilibrium populations of the PH and NPH were unequal, but the condition for mutant invasion was independent of host densities (see Section S2). When the pathogen had higher onward transmission on its PH (Case 3), by contrast, the equilibrium populations of each host type remained equal in size, and it was therefore unsurprising that there was no variation in ES virulence of their shared pathogen.

We further examined reciprocal host-host interactions where the interaction strengths were not equal, as this is more relevant for ecological scenarios where the interspecific interaction are unlikely to be the same between two species (Figure 5). As previously stated, under Case 1: Susceptibility specialism, the ES virulence in independent of host interactions (Figure 5A). For Case 2: Growth Rate specialism, ES virulence decreases as the NPH becomes a weaker competitor against the PH or a more generous mutualist to the PH (Figure 5B). Conversely, for Case 3: Onward Transmission and Case 4: Death Rate specialism ES virulence increases as the NPH become a weaker competitor or a more generous mutualist to the PH. For Case 3, the ranges of ES virulence for unequal competition or mutualism overlap with one another. For example, the scenario in which the NPH is a stronger competitor than the PH will actually result in a parasite with lower ES virulence than the scenario in which the NPH is a less-generous mutualist (see supplemental Figure S4.2 for absolute ES virulence values). This is unlike reciprocal, unequal strength relationships under Case 2 and Case 4 specialism (Fig. 5B, D). For these, mutualistic relationships, even unbalanced ones, tended to result in lower parasite virulence than competitive ones. Thus, in Case 3: Onward Transmission specialism, it is the relative relationship between hosts, rather than absolute sign, that is most significant in predicting shared pathogen virulence.

**Figure 5:**
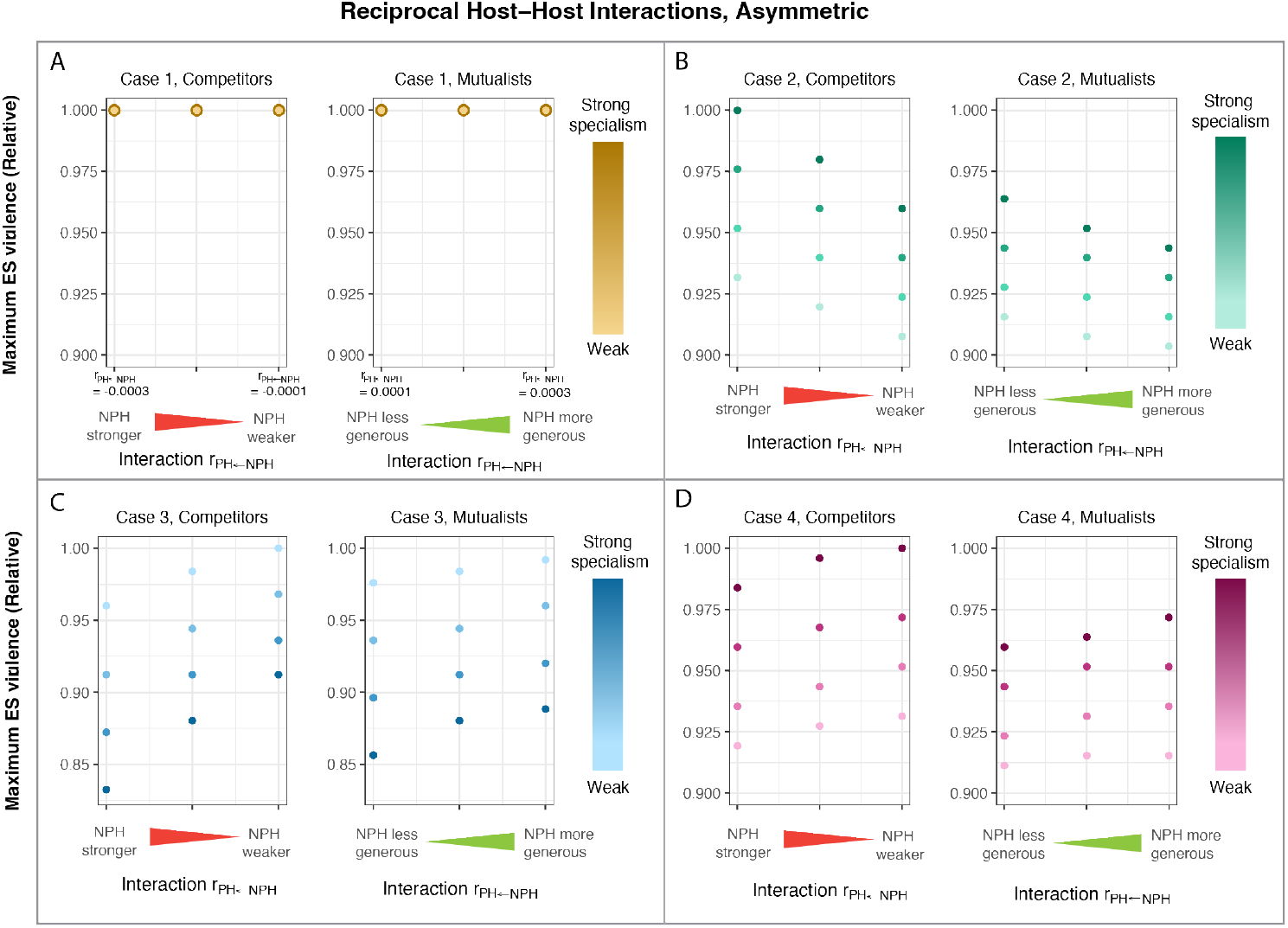
ES virulence for parasites of two reciprocally, but asymmetrically, interacting hosts. In each of the interactions tested here, the host interaction parameters had the same sign. The impact of the preferred host (PH) on the non-preferred host (NPH) was held constant (*r*_*NPH*←*PH*_ = 0.0002 for the mutualism panels, and *r*_*NPH*←*PH*_ = −0.0002 for the competition panels). The horizontal axes then indicate the strength of the impact of the non-preferred host on the preferred host (0.0001 < *r*_*P*_ |_*H*←*NPH*_ | < 0.0003; illustrated by the wedges), with resulting imbalances in the inter-host relationship noted. For example, when the NPH has a larger positive (negative) impact on the PH than the positive (negative) impact of the PH on the NPH, the NPH is necessarily a more generous mutualist (stronger competitor). Exact tick mark values are noted for the Case 1 panels, and are the same for all other Cases. We tested the host-host interactions on the backdrop of four different mechanisms of parasite specialism (Cases 1-4, panels A-D, respectively) and for four degrees of parasite specialism (ranging from strong to weak) per mechanism. The maximum ES virulence refers to the greater realized virulence, when realized virulence differs between host types (as occurs for Case 2 and 4 specialism). For the Case 1 plot, the degree of parasite specialism did not impact the ES virulence thus each point reflects four that are superimposed. For the exact parameter values for the degree of specialism, see Section S3.

### Non-reciprocal interactions

In a non-reciprocal (+/-) host-host interaction, knowledge of the mechanism of parasite specialism was necessary to predict the direction of virulence evolution. For Case 3, in which onward transmission was more efficient from the PH or in Case 4, in which excess death was reduced in the PH, higher virulence evolved when the exploiter was the PH and the exploitee was the NPH (Figure 6C,D). In contrast, for Case 2, in which the parasite grew more rapidly in the PH, higher virulence evolved when the exploitee was the PH and the exploiter was the NPH (Figure 6B). Figure 6 shows results for equal strength, non-reciprocal interactions, but the trends also hold for unequal strength, non-reciprocal interactions (see Figure S4.4). The case in which susceptibility differed between hosts once again did not show any shifts in ES virulence (Figure 6A).

**Figure 6:**
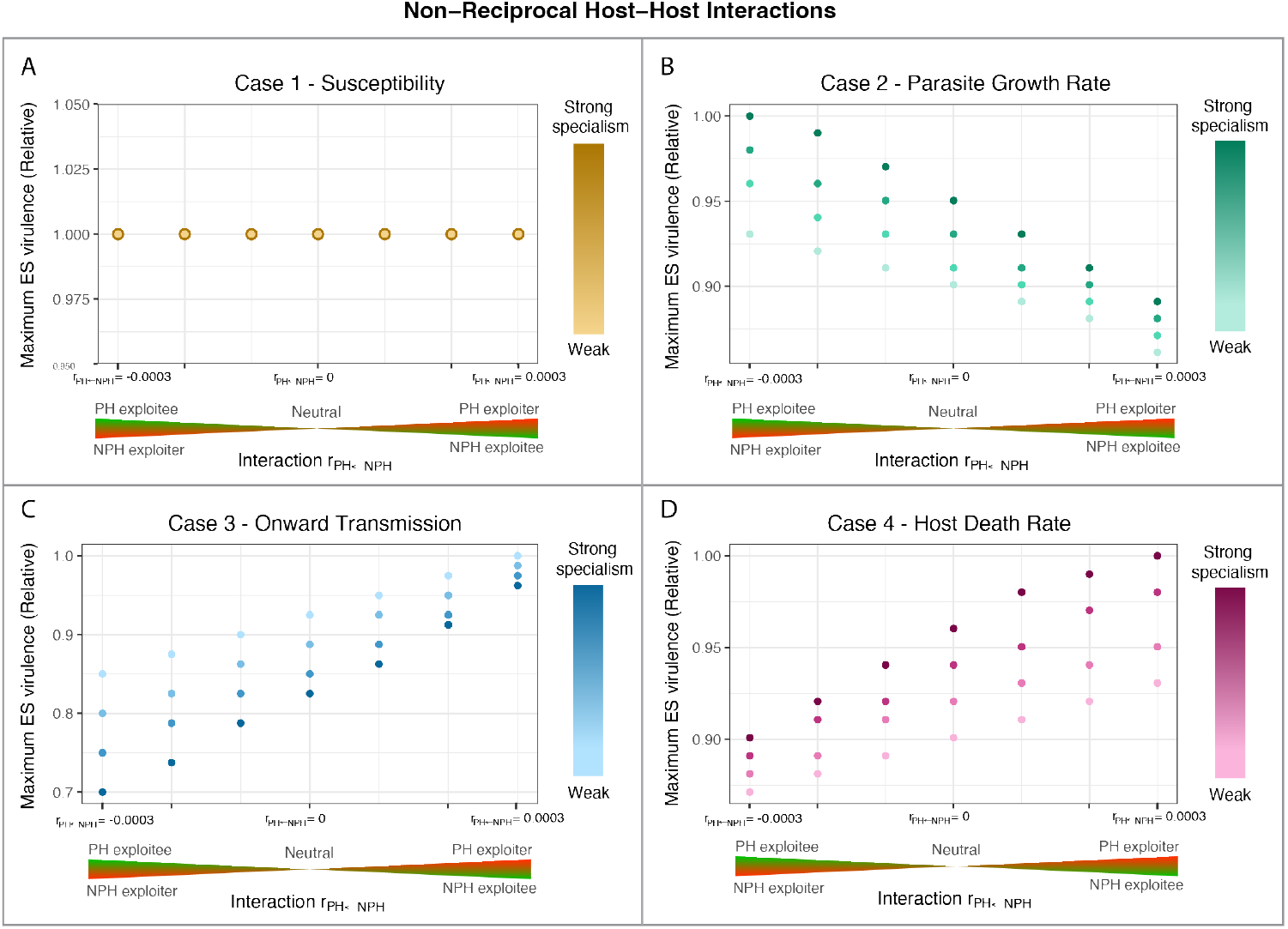
ES virulence for parasites of two non-reciprocally interacting hosts. In all the interactions tested here the two host interaction parameters had opposing signs: *r*_*ij*_ = −*r*_*ji*_. The horizontal axis indicates the impact of the non-preferred host on the preferred host. The left half of each Case panel, for example, represents interactions in which the non-preferred host exploits the preferred host: *r*_*PH*←*NPH*_ < 0 (red wedge) and *r*_*NPH*←*PH*_ > 0 (green wedge). We tested host-host interactions ranging from neutrality (|*r*_*ij*_| = |*r*_*ji*_| = 0) to strong exploitation (|*r*_*ij*_| = |*r*_*ji*_| = 0.0003). We tested these interactions on the backdrop of four different mechanisms of parasite specialism (Cases 1-4, panels A-D, respectively) and for four degrees of parasite specialism (ranging from strong to weak) per mechanism. The maximum ES virulence refers to the greater realized virulence, when realized virulence differs between host types (as occurs for Case 2 and 4 specialism). Because we do not compare magnitudes of ES virulence between cases, only within cases, we plot relative ES virulence (scaled from the highest value for a given specialism case). For the Case 1 plot, the degree of parasite specialism did not impact the ES virulence thus each point reflects four that are superimposed. For the exact parameter values for the degree of specialism, see Section S3.

When comparing the absolute ES virulence values between the reciprocal and non-reciprocal host relationships, the ‘extremes’ of parasite virulence were selected for in non-reciprocal host relationship scenarios (see Figures S4.1-4 for absolute ES virulence values). For example, in Case 2: Growth Rate specialism, the virulence selected for when the PH is the exploiter was *lower* than virulence selected for when the PH and NPH are mutualists, and the virulence when the PH is exploited is *higher* than the virulence when the PH and NPH are competitors.

### Impacts of the strength of parasite specialism

In addition to examining the role of host-host interactions, we assessed the interaction of host-host interactions with the strength of specialism (Figures 4-6). Regardless of whether we considered reciprocal or non-reciprocal relationships of equal or unequal strengths, Case 2: Growth Rate and Case 4: Death Rate specialism saw the evolution of higher virulence as disparities between parasite performance on its PH vs NPH grew stronger. In contrast – with similar interaction types and strengths of interaction – Case 3: Onward Transmission specialism showed the opposite trend: stronger specialization on the PH resulted in the evolution of lower virulence.

## Discussion

With a suite of models we have demonstrated the importance of the community context of multiple hosts with different interspecific interactions to the evolution of pathogen virulence. We have shown that parasite specialism is critical to the outcome, with the evolution of purely generalist parasites unaffected by the nature of the ecological interaction between their hosts. Furthermore, the strength of specialism has a direct impact on the evolutionarily stable (ES) virulence, alongside the degree of asymmetry of the host interaction and the fundamental nature of the interaction. Taken as a whole our work emphasises the importance of species interactions in determining the evolution of parasite virulence, and moreover emphasizes the community context of evolutionary outcomes. Moving beyond the simple host-parasite evolutionary models to include more realistic multi-species interactions is therefore an important aim for understanding the drivers of virulence in nature.

We can summarize the key impacts on virulence evolution in a shared parasite system as three fundamental factors: the strength of specialism on each host, the relative ecological relationship between hosts, and whether relationships could be classified as competitive or mutualistic. First, increasing the strength of specialism monotonically shifted ES virulence, regardless of hosthost interactions. Second, extremes of virulence were evolved in contexts where hosts had a large relative difference in their impacts on one another – for example, in cases of exploitation that was strongly beneficial for the exploiter and strongly detrimental to the exploitee. Third, in general, mutualistic relationships (even unbalanced ones) selected for lower shared parasite virulence than competitive relationships. Along with these commonalities, different mechanisms of parasite specialism differed in the direction of the relationship between strength of specialism and virulence, and in the direction of virulence evolution as we moved from a scenario where a preferred host was exploited to where it was the exploiter. Here we explain how of the outcomes of each case tie back to a unifying principle, most clearly illustrated by the exploiter/exploitee host-host relationship.

In general, the parasite/pathogen is competing with the exploiter for the exploitee resource, so it will evolve to be able to consume that resource more rapidly (and, thus, will cause more damage to the host) when the exploitee resource is relatively more threatened. We will break this trend down case by case. The case of reduced excess death on the preferred host (Case 4) is the most straightforward. Higher virulence evolves when the exploiter is the preferred host because the exploitee, as the non-preferred host, thus has higher disease-induced death. The case of more rapid growth in the preferred host (Case 2) is also fairly intuitive. Higher virulence evolves when the exploitee is the preferred host, since, due to the trade-off between parasite growth rate and virulence, higher parasite growth rate in the exploitee will directly lead to higher excess death.

The case in which there is increased onward transmission from the preferred host (Case 3) is slightly more complex, and requires consideration of feedback between parasite generations. Importantly, hosts that share ‘Case 3’ parasites share the same burden of infection (i.e. the same susceptiblity to infection from an existing pool of parasites and the same likelihood of death from infection). They differ only in how heavily their infection contributes to the common pool of parasite propagules. When, for example, a shared parasite produces more propagules for transmission on the exploitee, the exploiter’s negative impacts on the exploitee lessens relative competition for the exploitee resource, because the parasite will have lower reproduction rates on the more-available, non-preferred exploiter host. In the opposite case, exploitation strengthens competition, because the parasite has a higher reproduction rate on the more-available, preferred exploiting host. Thus, higher virulence evolves when the exploiter is the preferred host. These same trends from explicitly non-reciprocal host relationships hold when considering imbalanced, yet reciprocal, relationships. For example, consider a mutualistic host-host relationship with Case 2: Growth Rate specialism (Figure 5B). In this scenario, higher virulence evolves when the preferred host is more generous (i.e. closer to an exploitee than an exploiter) than when it is less generous than its mutualistic partner. Furthermore, this principle can explain commonalities in the observed trends along the competition to mutualism axis in Case 2 and Case 4: Death Rate specialism. In competition, hosts reduce one anothers’ abundance, but with less direct benefit to themselves (as compared to a direct exploiter). Thus, the parasite of two competitors still faces a more restricted host resource than does a parasite of two mutualists.

The relationship between the strength of specialism and the evolution of virulence is similarly explained by competition for hosts – if stronger specialism directly exacerbates the competition for quality hosts, higher virulence will evolve, and vice versa. In general, stronger specialism results in a reduction in the average quality of the host pool (for example, consider the extreme case in which a parasite can only infect one of the two hosts – this parasite would see the pool of the two types as inferior to a parasite who could equally parasitize both). In Cases 2: Growth Rate and 4: Death Rate specialism, this reduction in ‘average’ host quality directly strengthens competition for infecting the remaining pool of quality hosts, and thus higher virulence evolves when there is a strong difference in parasite performance on each host. Unlike in Case 2 or Case 4, stronger Case 3: Onward Transmission specialism does not restrict the number of hosts that will support rapid growth or longer infectious periods. For Case 3 specialism, we consider the consequences of infection to understand trends in ES virulence (the trends are reversed in Case 3 compared to Case 2 and 4) where it is *weaker* specialism that exacerbates competition for hosts. A virus that exhibits weaker (Case 3) specialism will generate relatively more propagules for onward transmission, and there will thus be more competition among viral lineages for seeding initial infections.

The consideration of host-host interactions in our model draws attention to scenarios in which previous work from one-host, one-parasite systems holds, and in which it does not. One interesting deviation is the role of host availability. High host availability is expected to select for higher virulence,^50^ and host scarcity for lower virulence. This is exemplified by the self-shading effect in spatial models, in which lower virulence is selected for when parasites would otherwise over-exploit their limited pool of neighboring hosts.^6, 51^ Once feedback of ecological relationships between multiple hosts is incorporated, however, this expectation does not appear to hold. In our model a mutualistic relationship supports higher host populations than does competition, but competition selects for higher ES virulence for the shared parasite.

There are also fruitful comparisons to draw to multi-host work that focused on intra-vs inter-host transmission and host life history parameters. Gandon^17^ established that parasites should evolve virulence that is optimal on their prime host (i.e. one that is present at high frequency and supports high parasite reproduction). When prime hosts are also more resistant, higher virulence is expected to evolve. Case 4 specialism, where the mechanism of specialism is a more resistant, tolerant host, demonstrates nicely how the impacts of host interactions on population dynamics can be incorporated into existing theory. In this case, parasite growth rates and reproduction are equivalent on both hosts, so class reproductive values will be primarily based on the relative proportion of hosts. Host-host relationships in which the more resistant PH is also ‘prime’ due to its higher frequency, such as in competition or as a direct exploiter, lead to the evolution of higher virulence (Figure 4D, 6D). Thus, we demonstrate that feedbacks from hosts’ ecological relationships can influence evolutionary trajectories of their shared parasites in ways previously achieved via a trade-off between resistance and host birth rate.^17^ However, it is not the full story; Case 3: Onward Transmission specialism shows that when we have a more complex network of host-host relationships, the effect may not hold for non-tolerance based definitions of resistance. In this case, higher virulence is selected for when the *less* resistant host (producing more propagules) is more abundant.

It is interesting to understand why Case 1 specialism, in which the parasite’s preferred host was more susceptible to initial infection than its non-preferred host, did not show dependence of ES virulence on either host-host relationships or on the degree of specialism. Mathematically (see Section S2), when we allowed a range of appropriate host populations to come to ecological equilibrium, their relative proportions were such that the condition for invasion (and thus, the ultimate ES virulence we would expect the system to evolve towards) was unchanged. Biologically, this model considers one-sided evolution of a parasite, and not co-evolution of the host – and Case 1 specialism is the only one of the four mechanisms that is independent of the parasite growth rate. Nevertheless, preliminary numerical simulations show that the time required for the invasion of a mutant parasite differs among competing, cooperating, and exploitative hosts (see Figure S5.1). There are times where the shared parasite system instead resembles a two-host, two-parasite system, which suggests that non-equilibrium parasite evolution in a community of hosts differing only in susceptibility is still an ecologically interesting system. In particular, pathogens (rabies, for example^52^) are known to produce variants that eventually become associated with distinct hosts.

In a world in flux, it is becoming more important than ever to characterize and predict the consequences of multi-host pathogens. Climate change,^53, 54^ deforestation and shifts in land usage,^55^ and a more interconnected globe^56, 57^ are all changing the strength and type of species interactions in existing communities and are exposing pathogens to potential new hosts. Hosts with shared parasites are at the forefront of of challenges in disease management. For example, disease-mediated invasions can reshape competition between native and invasive species,^3^ and zoonotic disease poses significant threats to human health.^58^ A combination of both future theory and empirical work is essential for predicting virulence evolution after spillover to new hosts, or in response to shifts in the host-host interactions themselves. Real ecological communities are, of course, more complex than the triads we study here, and it remains to be seen whether the principles carry over to larger host-parasite assemblages – though recent work in bottom-up predictions of community assembly^59^ are cause for hope that complicated ecological network processes can be predicted by considering subunits of the complex system. Furthermore, given that our work assumed a particular static mechanism of specialism, it would be interesting to examine the outcomes in a host-parasite coevolutionary context, where the degree of specialism and generalism in the parasite can evolve.^60^ Quantifying host-host interactions and the precise mechanisms of infection and transmission in a particular disease system is no small task, but it is our hope that integrating community ecology into epidemiological models will allow for better predictions of virulence evolution in a changing world.

## Acknowledgements

CE was supported by NSF-DGE-2146752. MB and AW were supported by the BBSRC/NSF EEID research grant BB/V00378X/1, NSF-DEB-2011109. MB was funded by NSF-DEB-2109860. This material is based upon work supported by the National Science Foundation Graduate Research Fellowship Program under Grant DGE 2146752. Any opinions, findings, and conclusions or recommendations expressed in this material are those of the author(s) and do not necessarily reflect the views of the National Science Foundation.

## Notes

### Competing Interest Statement

The authors have declared no competing interest.

