## Supplemental information for "Multispecies interactions and the community context of the evolution of virulence"

### Supplementary Information

#### S1 Deriving conditions for invasion of a rare mutant

To derive the invasion criteria of a mutant strain of the parasite we add the dynamics of the two hosts infected with the mutant strain $I_{1M}$ and $I_{2M}$ to the equations representing the resident 2-host, 1-parasite system. The model is represented as follows:

$$\frac{dS_1}{dt} = b(1-qN_1)S_1 - dS_1 + r_{12}N_2S_1 - \beta_{S1}\beta_{T1}S_1I_1 - \beta_{S1}\beta_{T2}S_1I_2 \tag{S.1}$$
$$- \beta_{S1M}\beta_{T1M}S_1I_{1M} - \beta_{S1M}\beta_{T2M}S_1I_{2M} + \gamma I_1 + \gamma I_{1M}$$
$$\frac{dI_1}{dt} = \beta_{S1}\beta_{T1}S_1I_1 + \beta_{S1}\beta_{T2}S_1I_2 - \gamma I_1 - (d+\alpha_1)I_1 \tag{S.2}$$
$$\frac{dS_2}{dt} = b(1-qN_2)S_2 - dS_2 + r_{21}N_1S_2 - \beta_{S2}\beta_{T2}S_2I_2 - \beta_{S2}\beta_{T1}S_2I_1 \tag{S.3}$$
$$- \beta_{S2M}\beta_{T2M}S_2I_{2M} - \beta_{S2M}\beta_{T1M}S_2I_{1M} + \gamma I_2 + \gamma I_{2M}$$
$$\frac{dI_2}{dt} = \beta_{S2}\beta_{T2}S_2I_2 + \beta_{S2}\beta_{T1}S_2I_1 - \gamma I_2 - (d+\alpha_2)I_2 \tag{S.4}$$
$$\frac{dI_{1M}}{dt} = \beta_{S1M}\beta_{T1M}S_1I_{1M} + \beta_{S1M}\beta_{T2M}S_1I_{2M} - \gamma I_{1M} - (d+\alpha_{1M})I_{1M} \tag{S.5}$$
$$\frac{dI_{2M}}{dt} = \beta_{S2M}\beta_{T2M}S_2I_{2M} + \beta_{S2M}\beta_{T1M}S_2I_{1M} - \gamma I_{2M} - (d+\alpha_{2M})I_{2M} \tag{S.6}$$

The parameters are as described in Table 1 in the main text. The addition of 'M' in a subscript indicates it is a parameter for the mutant (invading) parasite strategy. Note that superinfection is not possible in our model, and hosts do not have immunity or cross-immunity after they have recovered from an infection from either the wild-type or mutant parasite.

To determine the invasion criteria for the mutant parasite strategy we evaluation the Jacobian of the system of equations (S.1 - S.6) at the resident steady state where the mutant strain is absent (i.e. at $(\hat{S}_1, \hat{I}_1, \hat{S}_2, \hat{I}_2, 0, 0)$ ). Since we know the resident is at a stable steady state the invasion criteria for the mutant can be determined as the largest eigenvalue of $J_{mut}$ which is defined as follows:

$$J_{mut} = \begin{bmatrix} \frac{\partial \dot{I}_{1M}}{\partial I_{1M}} & \frac{\partial \dot{I}_{2M}}{\partial I_{1M}} \\ \frac{\partial \dot{I}_{2M}}{\partial I_{1M}} & \frac{\partial \dot{I}_{2M}}{\partial I_{2M}} \end{bmatrix},$$

and for our system,

$$J_{mut} = \begin{bmatrix} \beta_{S1M}\beta_{T1M}\hat{S}_1 - (d + \alpha_{1M} + \gamma) & \beta_{S1M}\beta_{T2M}\hat{S}_1 \\ \beta_{S2M}\beta_{T1M}\hat{S}_2 & \beta_{S2M}\beta_{T2M}\hat{S}_2 - (d + \alpha_{2M} + \gamma) \end{bmatrix}.$$

Solving for eigenvalues $\lambda$ gives us the condition

$$(\beta_{S1M}\beta_{T1M}\hat{S}_1 - (d + \alpha_{1M} + \gamma) - \lambda)(\beta_{S2M}\beta_{T2M}\hat{S}_2 - (d + \alpha_{2M} + \gamma) - \lambda) \\ -\beta_{S1M}\beta_{S2M}\beta_{T1M}\beta_{T2M}\hat{S}_1\hat{S}_2 = 0,$$

or, letting $\Gamma_{1M} = (d + \gamma + \alpha_{1M})$ and $\Gamma_{2M} = (d + \gamma + \alpha_{2M})$,

$$\lambda = \frac{1}{2}[B \pm \sqrt{B^2 - 4(-\Gamma_{1M}\beta_{S2M}\beta_{T2M}\hat{S}_2 - \Gamma_{2M}\beta_{S1M}\beta_{T1M}\hat{S}_1 + \Gamma_{1M}\Gamma_{2M}}),$$

where $B = \beta_{S2M}\beta_{T2M}\hat{S}_2 - \Gamma_{2M} + \beta_{S1M}\beta_{T1M}\hat{S}_1 - \Gamma_{1M}$. We can thus guarantee a real, positive eigenvalue ($\lambda_M$) if the mutant parasite invader meets the condition,

$$\lambda_M = \Gamma_{2M}\beta_{S1M}\beta_{T1M}\hat{S}_1 + \Gamma_{1M}\beta_{S2M}\beta_{T2M}\hat{S}_2 - \Gamma_{1M}\Gamma_{2M} > 0, \qquad \text{(S.7)}$$

or

$$\frac{\beta_{S1M}\beta_{T1M}\hat{S}_1}{\Gamma_{1M}} + \frac{\beta_{S2M}\beta_{T2M}\hat{S}_2}{\Gamma_{2M}} > 1. \qquad \text{(S.8)}$$

To use this fitness expression to determine the evolutionarily stable (ES) virulence of a particular host-host-parasite interaction, we first numerically allowed system of equations S.1 - S.6, with a resident parasite growth rate $\epsilon$, to equilibrate in the absence of a mutant parasite. The equilibrium values of $\hat{S}_1, \hat{S}_2$ were subsequently used in the fitness expression, with $\beta_{SiM}, \beta_{TiM}$ and $\Gamma_{iM}$ set by the mutant parasite's growth rate $\epsilon_M$ (see Section S.3 for exact values for each mechanism of parasite specialism), to determine whether invasion would occur. By plotting the binary outcome of whether $\lambda_M$ was positive or negative for each resident/mutant parasite combination, we generate Pairwise Invasibility Plots (PIPs) (as shown in Figure 3 in the main text).

#### S2 Independence of pure generalist parasites and Case 1 Specialism from host-host interactions

In the case where the mutant parasite is the same as the resident, the system will be in equilibrium and so the fitness expression, $\lambda_M = 0$. This is equivalent to the following expression:

$$\frac{\beta_{S1}\beta_{T1}\hat{S}_1}{\Gamma_1} + \frac{\beta_{S2}\beta_{T2}\hat{S}_2}{\Gamma_2} = 1. \tag{S.9}$$

**For a pure generalist parasite**, $\beta_{S1} = \beta_{S2}$, $\beta_{T1} = \beta_{T2}$, and $\Gamma_1 = \Gamma_2$. Thus, from Equation 9 the following will be true for the resident host-host-parasite equilibrium:

$$\frac{\beta_{S1}\beta_{T1}}{\Gamma_1}(\hat{S}_1 + \hat{S}_2) = 1. \tag{S.10}$$

For a mutant strain to invade requires

$$\frac{\beta_{S1M}\beta_{T1M}}{\Gamma_{1M}}(\hat{S}_1 + \hat{S}_2) > 1; \tag{S.11}$$

and so combining Equation S.10 and S.11 we can show that the mutant can invade if

$$\frac{\beta_{S1M}\beta_{T1M}}{\Gamma_{1M}} > \frac{\beta_{S1}\beta_{T1}}{\Gamma_1} \tag{S.12}$$

and the type that maximizes $\frac{\beta_{Si}\beta_{Ti}}{\Gamma_i}$ will win. These terms are independent of the $r_{ij}$ terms and of host densities. Therefore, host-host interactions will not impact the ES virulence of a pure generalist parasite.

**For Case 1 Specialism**, the preferred host (Host 1 for the following work) is more susceptible to initial infection than the non-preferred host. Thus, $\beta_{S1} > \beta_{S2}$, but $\beta_{S1} = \beta_{S1M}$, $\beta_{S2} = \beta_{S2M}$, $\beta_{T1} = \beta_{T2}$, and $\Gamma_1 = \Gamma_2$. From Equation 9 the following will be true for the resident host-host-parasite equilibrium:

$$\frac{\beta_{T1}}{\Gamma_1}(\beta_{S1}\hat{S}_1 + \beta_{S2}\hat{S}_2) = \frac{\beta_{T1}}{\Gamma_1}(\beta_{S1M}\hat{S}_1 + \beta_{S2M}\hat{S}_2) = 1. \tag{S.13}$$

The invasion condition becomes

$$\frac{\beta_{T1M}}{\Gamma_{1M}}(\beta_{S1M}\hat{S}_1 + \beta_{S2M}\hat{S}_2) > 1, \tag{S.14}$$

and combining equation S.13 and S.14 we can show that the mutant can invade if

$$\frac{\beta_{T1M}}{\Gamma_{1M}} > \frac{\beta_{T1}}{\Gamma_1} \tag{S.15}$$

. This condition does not depend on $r_{ij}$ terms or host densities, so the ESS virulence does not change as the host-host interactions change.

### S3 Parameter values for the numerical simulations

The following parameters were used in numerical simulations conducted in a portion of parameter space where the model's host-host-resident parasite equilibrium is stable, and both host types coexist (neither interspecific interactions nor infection drive either population to extinction). Full code for the numerical simulations can be found at https://github.com/cevensen/MultihostParasite.

| | Parameter(s) | Value |
|---|---|---|
| All simulations | $b_1, b_2$ | 1 |
| | $d_1, d_2$ | 0.1 |
| | $q_1, q_2$ | 0.0005 |
| | $\gamma_1, \gamma_2$ | 2 |
| | $r_{ij}$ | $0, \pm 0.0001, \pm 0.0002, \pm 0.0003$ |
| | $\beta_{11}, \beta_{12}, \beta_{22}, \beta_{21}$ | 0.1 |
| | $[\epsilon_{min}, \epsilon_{max}]$ | [3, 5.5] |

#### S3.1 Parameters for each mechanism of parasite specialism

| | Parameter(s) | Value |
|---|---|---|
| Case 1 Specialism | $\epsilon_1 = \epsilon_2 = \epsilon$ | Varied from $\epsilon_{min}$ to $\epsilon_{max}$ |
| | $\alpha_1 = \alpha_2$ | $\epsilon$ |
| | $\beta_{T1} = \beta_{T2}$ | $0.1\epsilon^{2/3}$ |
| | $\beta_{S1}$ | 0.1 |
| | $\beta_{S2}$ | [0.082, 0.084, 0.086, 0.088] |

The vector of values for $\beta_{S2}$ represent four different degrees of parasite specialism; 0.082 is the case with the highest degree of specialism.

| | Parameter(s) | Value |
|---|---|---|
| Case 2 Specialism | $\epsilon_1$ | Varied from $\epsilon_{min}$ to $\epsilon_{max}$ |
| | $\epsilon_2$ | $[0.82\epsilon_1, 0.84\epsilon_1, 0.86\epsilon_1, 0.88\epsilon_1]$ |
| | $\alpha_1$ | $\epsilon_1$ |
| | $\alpha_2$ | $\epsilon_2$ |
| | $\beta_{T1}$ | $0.1\epsilon_1^{2/3}$ |
| | $\beta_{T2}$ | $0.1\epsilon_2^{2/3}$ |
| | $\beta_{S1} = \beta_{S2}$ | 0.1 |

The vector of values for $\epsilon_2$ represent four different degrees of parasite specialism; $0.82\epsilon_1$ is the case with the highest degree of specialism.

| | Parameter(s) | Value |
|---|---|---|
| Case 3 Specialism | $\epsilon_1 = \epsilon_2 = \epsilon$ | Varied from $\epsilon_{min}$ to $\epsilon_{max}$ |
| | $\alpha_1 = \alpha_2$ | $\epsilon$ |
| | $\beta_{T1}$ | $0.1\epsilon^{2/3}$ |
| | $\beta_{T2}$ | $0.1\epsilon^{(2/3)-F}$ |
| | $F$ | [0.12, 0.10, 0.08, 0.06] |
| | $\beta_{S1} = \beta_{S2}$ | 0.1 |

The vector of values for $F$ represent four different degrees of parasite specialism; 0.12 is the case with the highest degree of specialism.

| | Parameter(s) | Value |
|---|---|---|
| Case 4 Specialism | $\epsilon_1 = \epsilon_2 = \epsilon$ | Varied from $\epsilon_{min}$ to $\epsilon_{max}$ |
| | $\alpha_1$ | $[0.82\epsilon, 0.84\epsilon, 0.86\epsilon, 0.88\epsilon]$ |
| | $\alpha_2$ | $\epsilon$ |
| | $\beta_{T1} = \beta_{T2}$ | $0.1\epsilon^{2/3}$ |
| | $\beta_{S1} = \beta_{S2}$ | 0.1 |

The vector of values for $\alpha_1$ represent four different degrees of parasite specialism; $0.82\epsilon$ is the case with the highest degree of specialism.

### S4 Absolute ES pathogen virulence

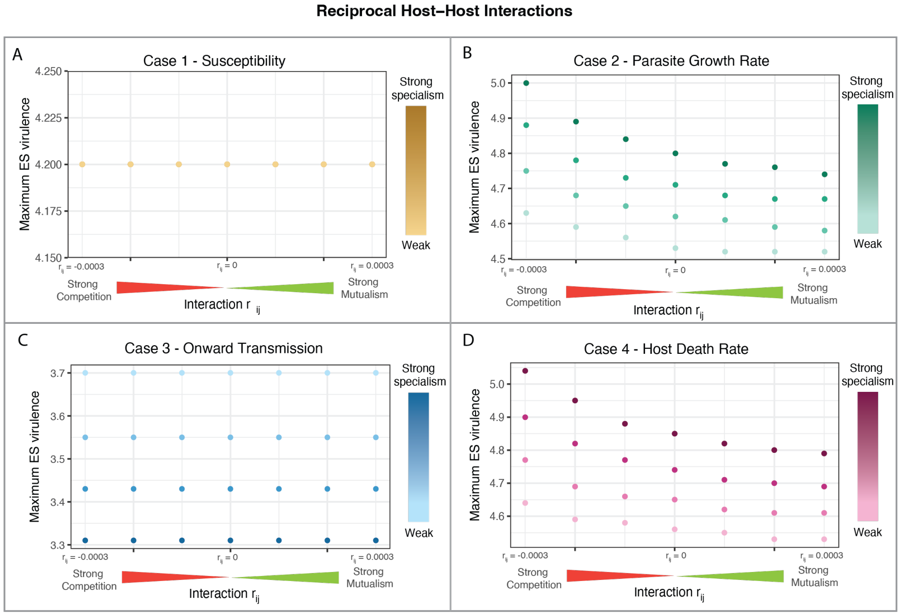


Figure S4.1: Absolute ES virulence for parasites of two reciprocally (and symmetrically) interacting hosts. In all the interactions tested here the two host interaction parameters were equal: $r_{ij} = r_{ji}$. Reciprocal host interactions ranged from strong competition ($r_{ij} = r_{ji} = -0.0003$) to strong mutualism ($r_{ij} = r_{ji} = 0.0003$). We tested the range of host interactions on the backdrop of four different mechanisms of parasite specialism (Cases 1-4, panels A-D, respectively). In addition, four degrees of parasite specialism (ranging from strong to weak) were tested for each mechanism – strong specialism meant there was a large difference in parasite performance on each host type. The maximum ES virulence (note vertical axes do not start at 0) refers to the greater realized virulence, when realized virulence differs between host types (as occurs for Case 2 and 4 specialism). For the Case 1 plot, the degree of parasite specialism did not impact the ES virulence thus each point actually reflects four that are superimposed. For the exact parameter values for the degree of specialism, see Section S3.

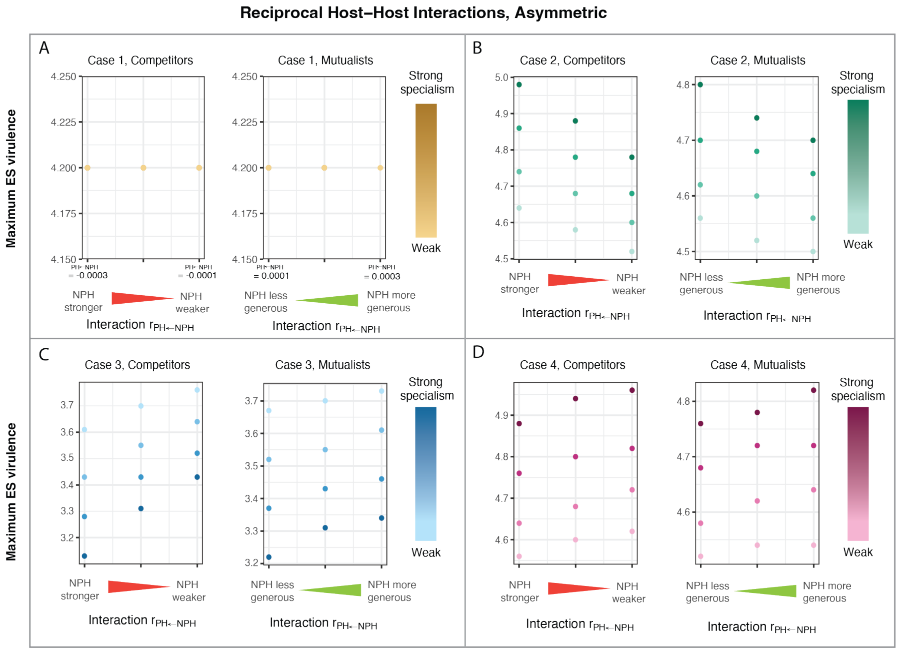


Figure S4.2: Absolute ES virulence for parasites of two reciprocally, but asymmetrically, interacting hosts. In each of the interactions tested here, the host interaction parameters had the same sign. The impact of the preferred host (PH) on the non-preferred host (NPH) was held constant ($r_{NPH\leftarrow PH} = 0.0002$ for the mutualism panels, and $r_{NPH\leftarrow PH} = -0.0002$ for the competition panels). The horizontal axes then indicate the strength of the impact of the non-preferred host on the preferred host ($0.0001 < |r_{PH\leftarrow NPH}| < 0.0003$; illustrated by the wedges), with resulting imbalances in the inter-host relationship noted. For example, when the NPH has a larger positive (negative) impact on the PH than the positive (negative) impact of the PH on the NPH, the NPH is necessarily a more generous mutualist (stronger competitor). Exact tick mark values are noted for the Case 1 panels, and are the same for all other Cases. We tested the host-host interactions on the backdrop of four different mechanisms of parasite specialism (Cases 1-4, panels A-D, respectively) and for four degrees of parasite specialism (ranging from strong to weak) per mechanism. The maximum ES virulence (note vertical axes do not start at 0) refers to the greater realized virulence, when realized virulence differs between host types (as occurs for Case 2 and 4 specialism). For the Case 1 plot, the degree of parasite specialism did not impact the ES virulence thus each point reflects four that are superimposed. For the exact parameter values for the degree of specialism, see Section S3.

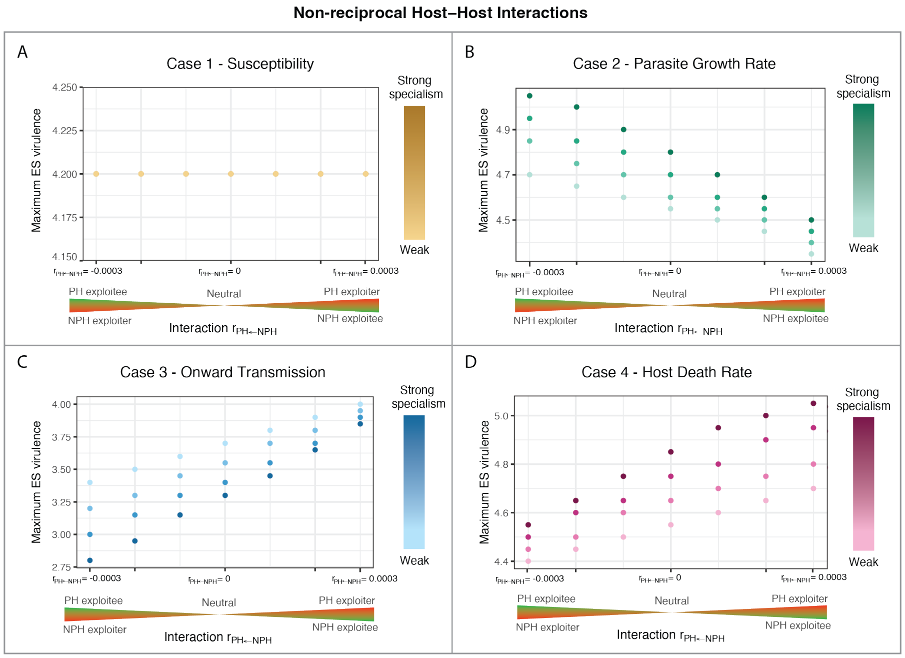


Figure S4.3: Absolute ES virulence for parasites of two non-reciprocally interacting hosts. In all the interactions tested here the two host interaction parameters had opposing signs: $r_{ij} = -r_{ji}$. The horizontal axis indicates the impact of the non-preferred host on the preferred host. The left half of each Case panel, for example, represents interactions in which the non-preferred host exploits the preferred host: $r_{PH \leftarrow NPH} < 0$ (red wedge) and $r_{NPH \leftarrow PH} > 0$ (green wedge). We tested host-host interactions ranging from neutrality ($|r_{ij}| = |r_{ji}| = 0$) to strong exploitation ($|r_{ij}| = |r_{ji}| = 0.0003$). We tested these interactions on the backdrop of four different mechanisms of parasite specialism (Cases 1-4, panels A-D, respectively) and for four degrees of parasite specialism (ranging from strong to weak) per mechanism. The maximum ES virulence (note vertical axes do not start at 0) refers to the greater realized virulence, when realized virulence differs between host types (as occurs for Case 2 and 4 specialism). For the Case 1 plot, the degree of parasite specialism did not impact the ES virulence thus each point reflects four that are superimposed. For the exact parameter values for the degree of specialism, see Section S3.

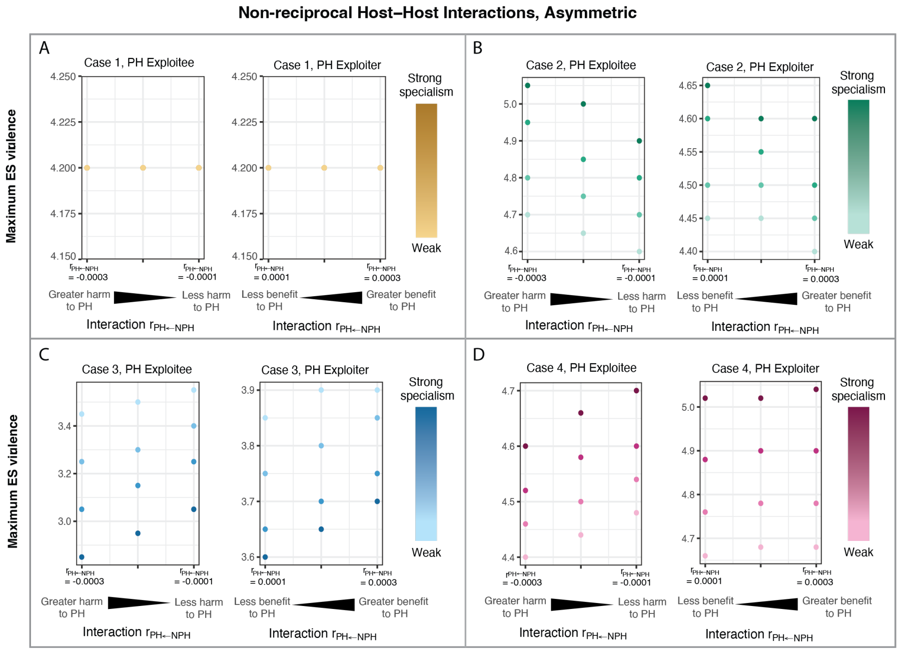


Figure S4.4: Absolute ES virulence for parasites of two non-reciprocally, asymmetrically, interacting hosts. In all the interactions tested here, the host interaction parameters had opposing signs. The impact of the preferred host (PH) on the non-preferred host (NPH) was held constant ($r_{NPH \leftarrow PH} = 0.0002$ for the PH Exploitee panels, and $r_{NPH \leftarrow PH} = -0.0002$ for the PH Exploiter panels). The horizontal axis indicates the impact of the non-preferred host on the preferred host. The PH Exploitee panels, for example, represent interactions in which the non-preferred host exploits the preferred host: $r_{PH \leftarrow NPH} < 0$ $(0.0001 < |r_{PH \leftarrow NPH}| < 0.0003)$ and $r_{NPH \leftarrow PH} = 0.0002$. The wedges indicate imbalances in the cost/benefit of exploitation. For example, the scenario in which the magnitude of the benefit the NPH receives by exploiting the PH ($|r_{NPH \leftarrow PH}|$) is larger than the magnitude of its negative impact on the PH ($|r_{PH \leftarrow NPH}|$) corresponds to the 'Less harm to PH' wedge label). We tested the host-host interactions on the backdrop of four different mechanisms of parasite specialism (Cases 1-4, panels A-D, respectively) and for four degrees of parasite specialism (ranging from strong to weak) per mechanism. The maximum ES virulence (note vertical axes do not start at 0) refers to the greater realized virulence, when realized virulence differs between host types (as occurs for Case 2 and 4 specialism). For the Case 1 plot, the degree of parasite specialism did not impact the ES virulence thus each point reflects four that are superimposed. For the exact parameter values for the degree of specialism, see Section S3.

#### S5 Transient invasion dynamics for Case 1 Parasite Specialism

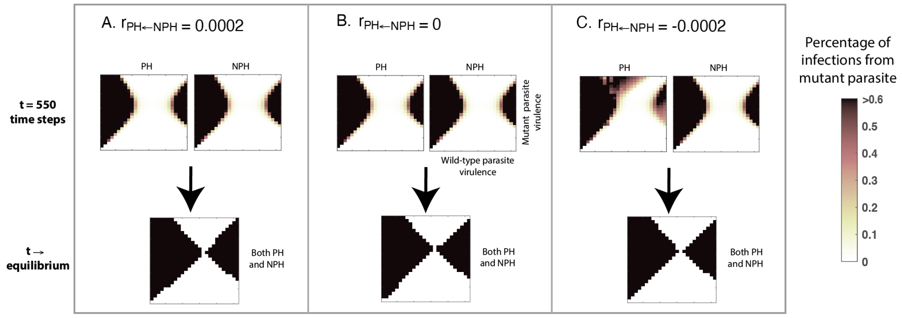


Figure S5.1. Status of attempted invasion by mutant parasite after 550 time steps (less than 10% of computational time typically needed to see stability), compared to the endpoint of an attempted invasion. The two hosts differ in initial susceptibility to the parasite (Case 1 specialism). In all panels, the preferred host (PH) has no impact on non-preferred host (NPH) (though we note the differences between panel C vs panels A and B are also seen in cases where the PH negatively or positively impacts the NPH). The attempted invasions are depicted very similarly to PIPs (see Figure 3 in the main text; wild-type or resident virulence is on the horizontal axis of each plot, and the invader's virulence is on the vertical axis), but here reflect the invasion dynamics in a quantitative manner. Darker regions reflect more active mutant infections in the PH (at left in each panel) or in the NPH (at right), approaching black when the mutant has displaced the resident and is responsible for 100% of infections from that point on. Though the attempted parasite invasions on each host may be distinct at early time points (a representative t=550 time steps is shown here), at equilibrium they eventually succeed (i.e. all infections, on both the PH and NPH, are due to the mutant) or fail (all the the infections on either host are from the original resident). As seen in panel C, when the less-susceptible host (NPH) negatively impacts the more-susceptible host (PH), higher virulence parasites can persist longer, resulting in some ultimately successful invasions occurring more slowly in the PH, and some ultimately unsuccessful invasions persisting longer in the PH. In the latter case, due to the invasion already having failed in the NPH, there exists an infection distribution akin to a two-host, two-parasite system.
